# The MAP kinase scaffold MORG1 shapes cell death in unresolved ER stress in Arabidopsis

**DOI:** 10.1101/2025.01.08.632046

**Authors:** Joo Yong Kim, Dae Kwan Ko, Federica Brandizzi

**Affiliations:** MSU-DOE Plant Research Lab, Michigan State University, East Lansing, Michigan, USA; Department of Plant Biology, Michigan State University, East Lansing, Michigan, USA; Great Lakes Bioenergy Research Center, Michigan State University, East Lansing, Michigan, USA

## Abstract

Governed by the unfolded protein response (UPR), the ability to counteract endoplasmic reticulum (ER) stress is critical for maintaining cellular homeostasis under adverse conditions. Unresolved ER stress leads to cell death through mechanisms that are yet not completely known. To identify key UPR effectors involved in unresolved ER stress, we performed an ethyl methanesulfonate (EMS) suppressor screen on the Arabidopsis *bzip28/60* mutant, which is impaired in activating cytoprotective UPR pathways. This screen identified MAP kinase organizer 1 (MORG1), a conserved MAP kinase scaffold protein, as a previously uncharacterized regulator of ER stress tolerance. The *coffin1* mutant, which carries a mutation in *MORG1*, exhibited enhanced resilience to ER stress by partially restoring UPR gene expression and promoting growth under stress conditions. Mechanistically, we found that MORG1 modulates MPK6-dependent phosphorylation of the stress-responsive transcription factor WRKY8. Loss of *WRKY8* phenocopied the *coffin1* mutant, highlighting WRKY8’s role as a key repressor in the UPR. Together, these findings reveal a MORG1–MPK6–WRKY8 signaling axis that fine-tunes UPR gene expression, providing new insights into ER stress regulation and strategies for improving stress tolerance in multicellular eukaryotes.

## Introduction

The endoplasmic reticulum (ER) is a central organelle in all eukaryotes, facilitating the synthesis and folding of proteins essential for cellular homeostasis and growth. Environmental stresses disrupt protein folding in the ER, leading to the accumulation of misfolded proteins, a condition referred to as ER stress^1^. To mitigate this, cells activate the unfolded protein response (UPR), a signaling network that restores ER homeostasis by enhancing the protein-folding capacity and promoting the degradation of misfolded proteins^1–3^.

The UPR is governed by ER stress sensors, which are highly conserved across taxa, the ribonuclease and kinase Inositol Requiring Enzyme 1 (IRE1) being the most conserved^1,4–6^. Multicellular eukaryotes have also evolved membrane-tethered transcription factors (TFs) as ER stress responsive effectors^7^. In plants, the UPR is primarily orchestrated by the TFs bZIP28 and bZIP60^2,3^. Under ER stress, bZIP28 is mobilized from the ER membrane to the Golgi apparatus where the cytosolic-facing TF domain is cleaved off the membrane and translocated to the nucleus^8,9^, while IRE1 splices the mRNA bZIP60 to generate its active form (sbZIP60)^10,11^. In the nucleus, bZIP28 and bZIP60 modulate the expression of overlapping and unique UPR target genes^12,13^. Loss of function in these TFs compromises UPR signaling, exacerbating plant sensitivity to ER stress^2,3^. These observations underscore the importance of tightly regulated ER stress responses for plant resilience.

While significant advances have been made in identifying the core components of the UPR, the broader regulatory framework governing ER stress responses remains incomplete. Recent studies have uncovered several key regulators that influence UPR signaling, either by acting downstream of bZIP28 and bZIP60 or by competing with them for binding to promoters of target genes. For example, the ubiquitin E3 ligase PIR1 acts downstream of IRE1 and negatively regulates the UPR by influencing the stability of the TF ABI5, a positive regulator of *bZIP60* expression^14^. Additionally, HY5, a positive regulator of light signaling, has been shown to antagonize the activity of bZIP28 and bZIP60 and repress UPR gene expression, highlighting the importance of transcriptional balancing in ER stress responses^15^. Furthermore, GBF2, a G-box binding TF, competes with bZIP28 and bZIP60 for binding to UPR target promoters, thereby repressing UPR gene expression and modulating the amplitude of the response^16^. These findings underscore the complexity of UPR regulation and suggest that a network of interacting factors fine-tunes the UPR signaling to ensure appropriate adaptation to ER stress. However, the full extent of this regulatory network and the precise mechanisms by which these factors interact remain to be fully elucidated. In particular, the roles of unconventional regulators, such as MAP kinase scaffolds and WRKY TFs, are largely unexplored in the context of ER stress. Further investigation into these regulatory mechanisms will be crucial for understanding the dynamic nature of the UPR and its role stress resilience. Indeed, emerging evidence suggests that MAP kinase signaling cascades^17–20^, which are central to various plant stress responses, may also influence the UPR^17^. Among these potential regulators is MORG1 (MAP kinase organizer 1), a conserved WD40 repeat protein known for its role as a scaffold in MAP kinase signaling across diverse organisms^21–24^. WRKY TFs are known to play diverse roles in plant stress responses, acting as both positive and negative regulators of gene expression depending on the specific context and interacting partners^25,26^. WRKYs are also emerging as potential regulators of the UPR. For example, some WRKY proteins have been shown to regulate the expression of UPR-related genes or interact with components of the UPR pathway^27,28^. Unraveling the molecular mechanisms underlying these signaling interactions, particularly the potential role of MORG1, could deepen our understanding of stress adaptation and inform strategies for engineering resilience in various organisms, including crops.

Among WRKY TFs, WRKY8 emerges as a key regulator involved in both biotic and abiotic stress responses. For example, previous studies showed that WRKY8 expression is induced by pathogen infection and wounding, and that it positively regulates resistance to *Botrytis cinerea* while negatively regulating resistance to *Pseudomonas syringae* ^29^. This suggests that WRKY8 plays a role in fine-tuning the balance between defense pathways. In addition to biotic stress, WRKY8 interacts with the VQ9 protein to modulate salinity stress tolerance, highlighting the involvement of protein-protein interactions in WRKY8-mediated stress responses^30^. Furthermore, WRKY8 promotes the transcriptional self-amplification of PBL19 and thereby boosts chitin-elicited, PBL19-EDS1-module-mediated basal immunity to *Verticillium dahliae*^31^. Interestingly, other members of the WRKY family, such as WRKY7, -11, and -17, negatively regulate the bZIP28-mediated UPR pathway during PAMP-triggered immunity^27^. These findings suggest that different WRKYs may play distinct or even opposing roles in modulating the UPR and other stress response pathways. Despite these advances, whether WRKY8 regulates the UPR and interacts with other signaling pathways, such as the MORG1-mediated MAP kinase signaling pathway, in the UPR remains largely unknown.

In this study, using *Arabidopsis thaliana* as a model species, we demonstrate that MORG1 is a key regulator of ER stress responses in Arabidopsis and define that its action pathway occurs via WRKY8’s ability to suppress UPR gene expression. Using an EMS suppressor screen in the ER-stress hypersensitive *bzip28/60* double mutant background, we identified a loss-of-function mutation in *MORG1* (*coffin1*) that enhances ER stress tolerance. Our study reveals the molecular mechanisms by which MORG1 regulates UPR gene expression via its interactions with MAP kinase signaling and WRKY8. These findings establish a MORG1–MPK6–WRKY8 signaling axis that fine-tunes UPR responses, providing novel insights into the regulatory landscape of ER stress and highlighting potential strategies for enhancing stress resilience in multicellular eukaryotes.

## Results

### Identification of *coffin1* as a suppressor of the lethality of *bzip28/60* to induced ER stress

To uncover critical regulators of the UPR acting downstream the essential UPR signaling axis exerted by bZIP28 and bZIP60, we performed an EMS suppressor screen in the hypersensitive *bzip28/60* double mutant background, which exhibits severe sensitivity to chronic ER stress induced by tunicamycin (TM), a specific ER stress inducer^32^. By screening approximately 20,000 M2 seedlings grown on plates supplemented by TM, we identified a mutant, *coffin1*, which exhibited significantly enhanced tolerance to TM-induced ER stress compared to *bzip28/60* (Figure 1a). Phenotypic analyses showed that *coffin1* plants displayed increased root elongation, higher fresh weight, and reduced ion leakage compared to *bzip28/60* mutants under TM treatment (Figure 1a–e). These results suggest that the *coffin1* mutation mitigates the effects of unresolved ER stress on cell survival in the absence of bZIP28 and bZIP60, implicating a novel pathway in ER stress resilience.

**Figure 1.**
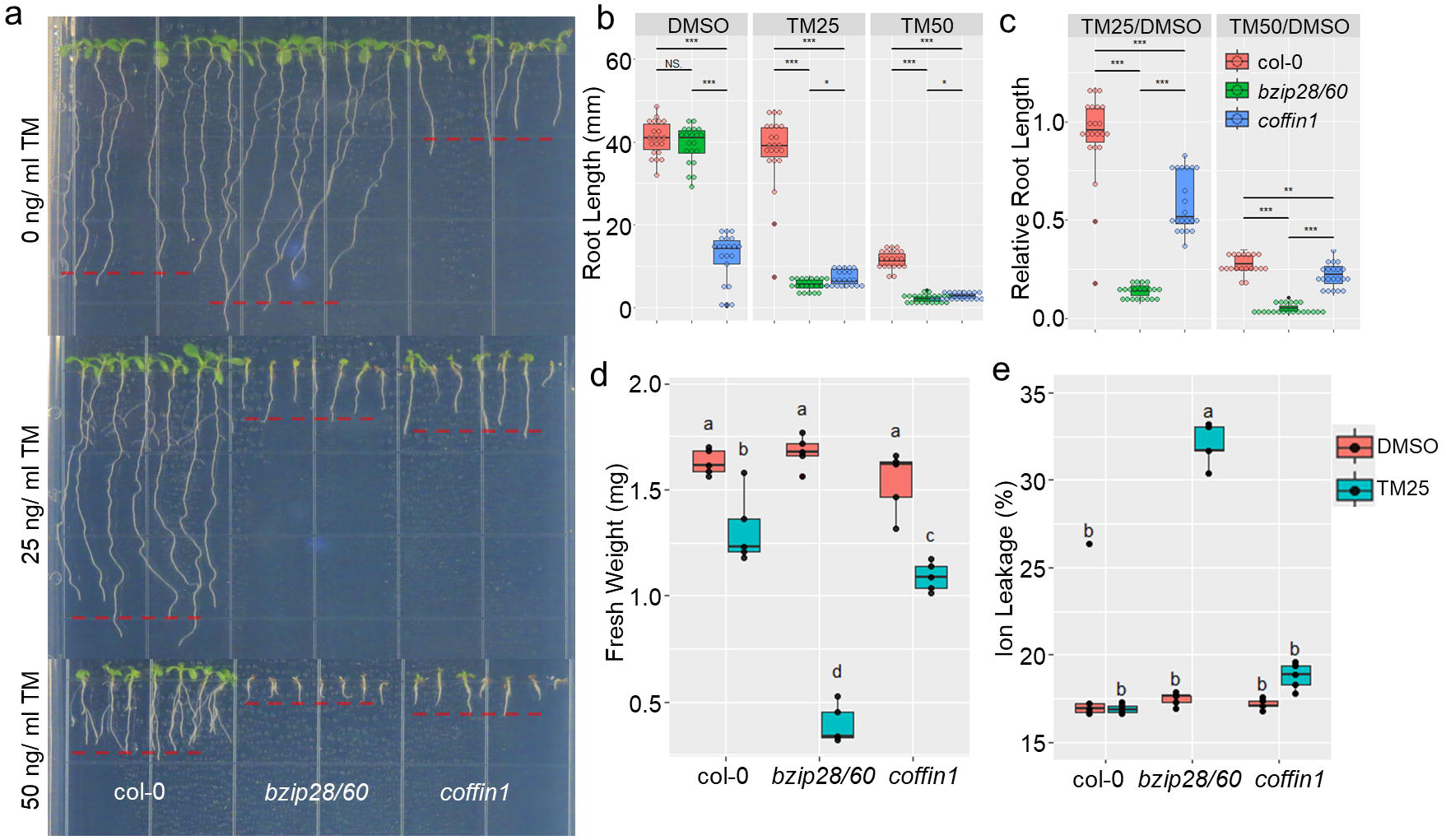
*coffin1* suppresses *bzip28/60* phenotype under chronic ER stress conditions. **a**. Phenotypic analysis of *coffin1* under chronic ER stress. seven-day-old seedlings of col-0, *bzip28/60*, and *coffin1* were grown on 1/2 LS media supplemented with DMSO (Mock), 25 ng/μl TM, or 50 ng/μl TM. Representative images of seedlings are shown. **b**. Root length of seedlings shown in **a**. Median ± IQR; *n* = 3 replicates with 7 seedlings per replicate. Statistical significance was determined by Student’s *t*-test. **c**. Relative root length of seedlings shown in **a**. Median ± IQR; *n* = 3 replicates with 7 seedlings per replicate. Statistical significance was determined by Student’s *t*-test. **d**. Fresh weight of seven-day-old col-0, *bzip28/60*, and *coffin1* seedlings grown on 1/2 LS media supplemented with DMSO or 25 ng/μl TM. Median ± IQR; *n* = 3 replicates with 7 seedlings per replicate. Statistical significance was determined by one-way ANOVA with Tukey’s post-hoc test. **e**. Ion leakage of seven-day-old col-0, *bzip28/60*, and *coffin1* seedlings grown on 1/2 LS media supplemented with DMSO or 25 ng/μl TM. Median ± IQR; *n* = 4 replicates. Statistical significance was determined by Student’s *t*-test.

### Mapping *coffin1* identifies the causal mutation in *MORG1*

To determine the genetic basis of the *coffin1* phenotype, we conducted bulk segregant analysis combined with whole-genome sequencing (WGS)^33,34^. The *coffin1* mutant (*bzip28/60/coffin1*) was crossed with *bzip28/60* to generate an F₂ mapping population segregating for the suppressor mutation. DNA was extracted from pooled individuals exhibiting the *coffin1* phenotype and those showing the *bzip28/60* phenotype under TM treatment (Figure S1a).

Analysis of single nucleotide polymorphism (SNP) frequency across the genome revealed a prominent peak on chromosome 5, identifying a candidate region harboring the causal mutation (Figures 2a, S1b). Within this region, we found a missense mutation in the *MORG1* gene (*AT5G64730*), resulting in a putative serine-to-phenylalanine substitution at position 288 (S288F) within the conserved WD40 repeat domain of the protein (Figures 2b, S1c).

**Figure 2.**
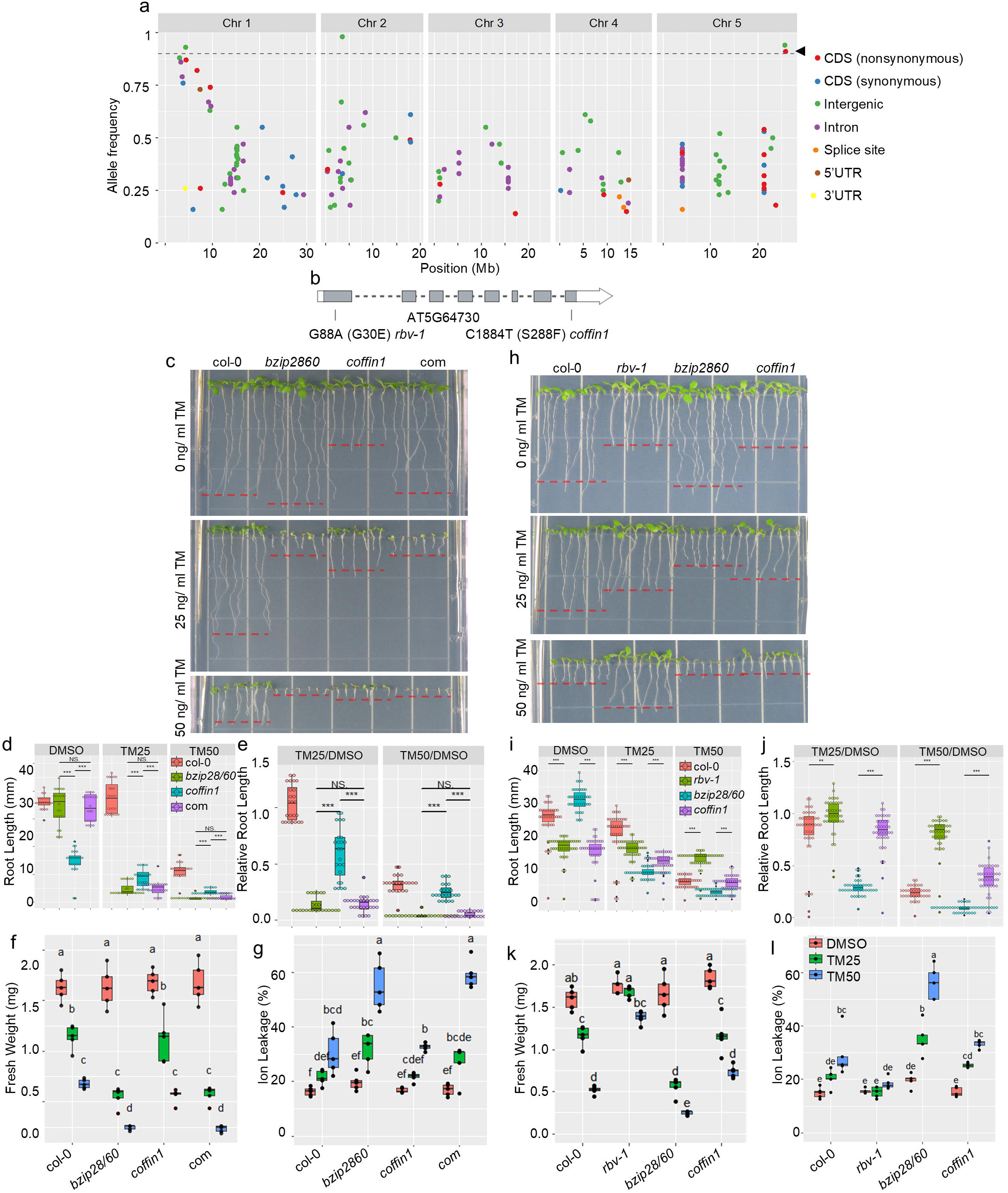
MORG1 is a causal mutation of *coffin1*. **a.** Allele frequencies of single nucleotide polymorphisms (SNPs) in the ER stress-resistant pool of *coffin1* BC1F2. Different colors indicate different different types of SNPs, as shown on the bottom of the plot. The dashed line indicate 0.9 allele frequency. The back arrowhead indicates the non-synonymous mutation in the CDS of *AT5G64730 (MORG1)*. **b**. Schematic diagram of the *MORG1*, highlighting the locations of point mutations in *coffin1* and *rbv-1*. **c**. Phenotypic analysis of *coffin1* complemented with MORG1 expression. *Coffin1* and *coffin1* complemented with MORG1 were grown on 1/2 LS media supplemented with DMSO (Mock) or 25 ng/μl TM. Representative images are shown. **d**. Root length of seedlings shown in **c**. Median ± IQR; *n* = 3 replicates with 7 seedlings per replicate. Statistical significance was determined by Student’s *t*-test. **e**. Relative root length of seedlings shown in **c**. Median ± IQR; *n* = 3 replicates with 7 seedlings per replicate. Statistical significance was determined by Student’s *t*-test. **f**. Fresh weight of seedlings shown in **c**. Median ± IQR; *n* = 4 replicates (8 seedlings pooled per replicate). Statistical significance was determined by one-way ANOVA with Tukey’s post-hoc test. **g**. Ion leakage of seedlings shown in **c**. Median ± IQR; *n* = 5 replicates (10 seedlings pooled per replicate). Statistical significance was determined by one-way ANOVA with Tukey’s post-hoc test. **h**. Phenotypic analysis of col-0, *rbv-1*, *bzip28/60*, and *coffin1* under chronic ER stress. Seven-day-old seedlings were grown on 1/2 LS media supplemented with DMSO (Mock), 25 ng/μl TM, or 50 ng/μl TM. Representative images are shown. **i**. Root length of seedlings shown in **h**. Median ± IQR; *n* = 3 replicates with 7 seedlings per replicate. Statistical significance was determined by Student’s *t*-test. **j**. Relative root length of seedlings shown in **h**. Median ± IQR; *n* = 3 replicates with 7 seedlings per replicate. Statistical significance was determined by Student’s *t*-test. **k**. Fresh weight of seedlings shown in **h**. Median ± IQR; *n* = 5 replicates (7 seedlings pooled per replicate). Statistical significance was determined by one-way ANOVA with Tukey’s post-hoc test. **l**. Ion leakage of seedlings shown in **h**. Median ± IQR; *n* = 4 replicates (10 seedlings pooled per replicate). Statistical significance was determined by one-way ANOVA with Tukey’s post-hoc test.

*MORG1* encodes MAP kinase organizer 1, a scaffold protein that facilitates MAP kinase signaling cascades^23^. To confirm that the mutation in *MORG1* was responsible for the *coffin1* phenotype, we introduced a wild-type *MORG1* CDS driven by its native promoter into *coffin1* plants. Complementation fully restored the ER stress-sensitive phenotype, resembling the parental *bzip28/60* mutant (Figure 2c–g), thereby validating the causal role of the *MORG1* mutation in conferring ER stress tolerance.

### Loss of MORG1 function confers ER stress resistance

To investigate whether the loss of *MORG1* function alone enhances ER stress tolerance, we analyzed a previously characterized *morg1* allele, *rbv-1* (*REDUCTION IN BLEACHED VEIN AREA*), which carries a G88A (G30E) point mutation in the MORG1 protein in col-0 background^35^. Similar to *coffin1*, the *rbv-1* mutant exhibited increased root elongation, higher fresh weight, and reduced ion leakage compared to WT under chronic TM-induced ER stress (Figure 2h-l). These results demonstrate that loss of MORG1 function enhances ER stress tolerance not only in the *bzip28/60* background but also in a background where bZIP28 and bZIP60 are present, highlighting a master regulatory role of MORG1 in regulating cell fate.

### Transcriptome analyses reveal partial restoration of UPR gene expression in *coffin1*

To elucidate the molecular basis of enhanced ER stress tolerance in *coffin1*, we conducted RNA-seq on WT, *bzip28/60*, and *coffin1* seedlings treated with DMSO (control) or 500 ng/mL TM for 6 hours. Initial analysis using a pairwise Pearson correlation plot revealed high similarity between biological replicates and distinct clustering of samples based on genotype, with *coffin1* showing the most distinct transcriptomic profile (Figure 3a). This confirms the reliability of our transcriptome profiling and suggests substantial transcriptomic changes in *coffin1*.

**Figure 3.**
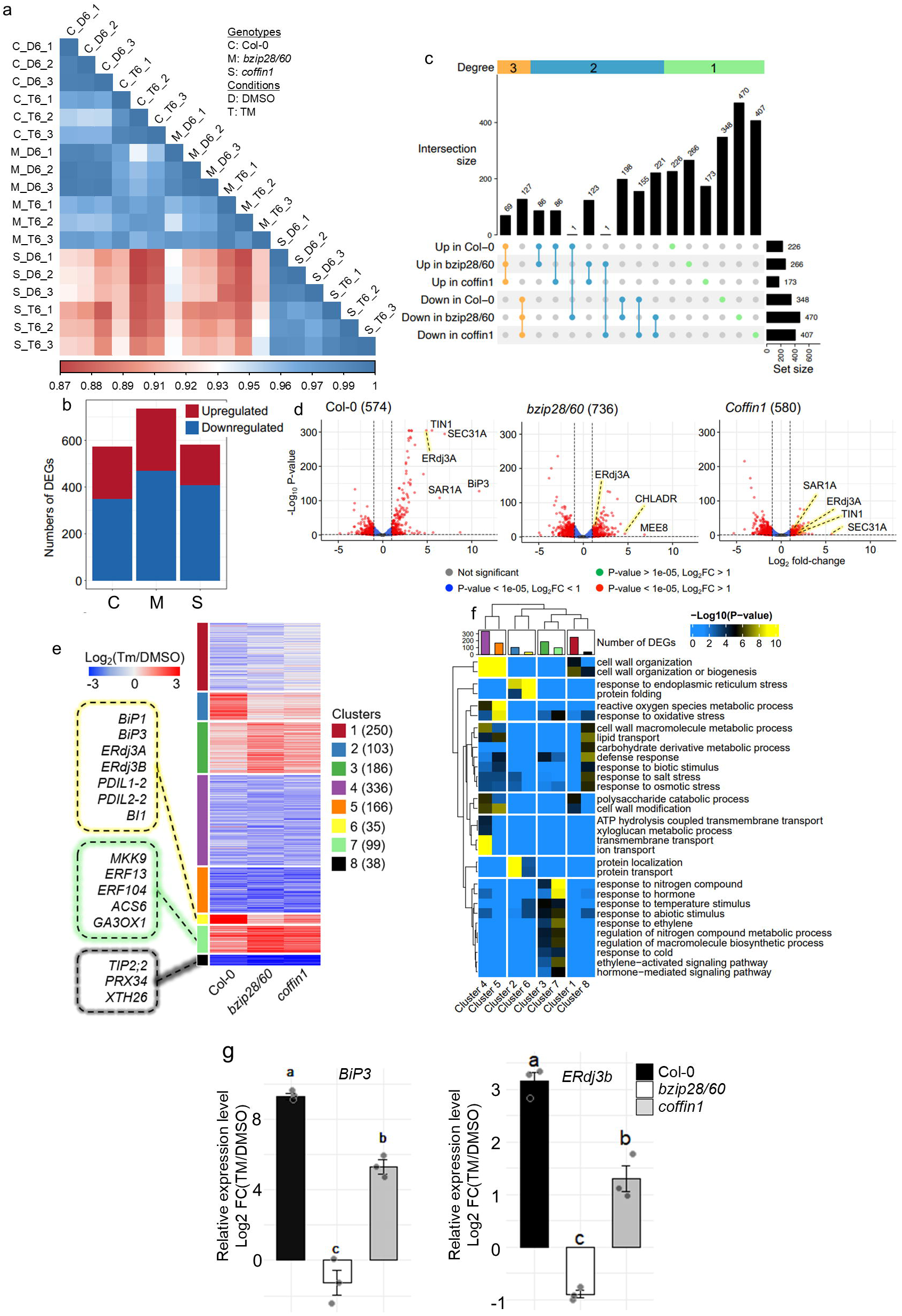
Transcriptome analysis reveals partial restoration of UPR gene expression in *coffin1*. **a.** Pairwise Pearson correlation plot of RNA-seq data from col-0, *bzip28/60*, and *coffin1* seedlings treated with DMSO (mock) or 500 ng/mL TM for 6 hours. The color scale represents the Pearson correlation coefficient (r), with blue indicating high correlation and red indicating low correlation. **b**. Number of differentially expressed genes (DEGs) in col-0, *bzip28/60*, and *coffin1* seedlings treated with 500 ng/mL TM for 6 hours compared to their respective DMSO (mock) controls. **c**. Upset plot showing the intersections of DEGs across genotypes. Each bar represents the number of DEGs in each intersection set. **d**. Volcano plots showing differentially expressed genes in col-0, *bzip28/60*, and *coffin1* seedlings treated with 500 ng/mL TM for 6 hours compared to their respective DMSO (mock) controls. The x-axis represents the log2 fold change (log2FC) of gene expression in TM relative to mock. The y-axis represents the -log10 (adjusted p-value). Red dots indicate differentially expressed genes, blue dots indicate genes not significantly differentially expressed. Selected UPR genes are highlighted. **e**. K-means clustering analysis of differentially expressed treated with 500 ng/mL TM for 6 hours compared to their respective DMSO (mock) controls. Eight clusters were identified based on their expression patterns. The heatmap shows the normalized expression values of each gene in each sample. Clusters 6, 7, and 8, which show the most pronounced differences between *coffin1* and *bzip28/60*, are highlighted. **f**. Gene Ontology (GO) enrichment analysis of DEGs by clusters in **e**. **g**. Validation of RNA-seq data by quantitative real-time PCR (qRT-PCR). The relative expression levels of *BiP3* and *ERdj3B* were measured in col-0, *bzip28/60*, and *coffin1* seedlings treated with DMSO (mock) or 500 ng/mL TM for 6 hours. Data are normalized to UBQ10 and presented as means ± SD (n = 3 biological replicates).

To identify potential functional targets of MORG1, we analyzed the transcriptomes of WT, *bzip28/60*, and *coffin1* genotypes to pinpoint differentially expressed genes (DEGs) under treatment conditions for each genotype. In response to TM, we identified 348 downregulated and 226 upregulated genes in WT, 470 downregulated and 266 upregulated genes in *bzip28/60*, and 407 downregulated and 173 upregulated genes in *coffin1* (Figure 3b). An upset plot further illustrated the relationships between these DEG sets, highlighting both unique and overlapping gene expression changes across genotypes (Figure 3c). Next, we focused on identifying genes with distinct expression patterns between *bzip28/60* and *coffin1* under TM treatment, as these genes are likely to play a role in the *coffin1* phenotype. Volcano plot analysis revealed that a subset of UPR genes was induced by TM in *coffin1* but not in *bzip28/60* (Figure 3d), suggesting a partial restoration of UPR gene expression in *coffin1*. To further characterize these differentially expressed genes, we performed k-means clustering analysis, which grouped them into eight distinct clusters based on their expression patterns (Figure 3e). We focused on clusters 6, 7, and 8, as these clusters exhibited the most pronounced differences between *bzip28/60* and *coffin1*. Cluster 6 (35 genes) was particularly interesting, as it contained a significant number of UPR genes that were highly induced in WT, not induced in *bzip28/60*, and partially induced in *coffin1* under TM treatment. A Gene Ontology (GO) enrichment analysis of cluster 6 further confirmed the overrepresentation of UPR-related terms, including “response to endoplasmic reticulum stress” and “protein folding” (Figure 3f).

To validate the RNA-seq data and confirm the partial restoration of UPR gene expression, we performed quantitative real-time PCR (qRT-PCR) on selected UPR genes, including *BiP3* and *ERdj3B*. Consistent with the RNA-seq results, qRT-PCR analysis confirmed that the induction of these genes was partially restored in *coffin1* compared to *bzip28/60* under TM treatment (Figure 3g).

Taken together, these results suggest that the MORG1 mutation in *coffin1* facilitates the partial reactivation of specific UPR pathways, compensating for the complete loss of bZIP28 and bZIP60 function. The identification of UPR genes in cluster 6, which are partially induced in *coffin1* but not in *bzip28/60*, provides a crucial link between MORG1 function and UPR gene regulation and sets the stage for further investigation into the underlying molecular mechanisms.

### Promoter analysis suggests involvement of NAC and WRKY binding motifs

To identify potential TFs responsible for the partial restoration of UPR gene expression in *coffin1*, we conducted a promoter analysis of the DEGs identified in our RNA-seq analysis, focusing on genes that showed increased expression in *coffin1* compared to *bzip28/60* under TM treatment (Figure 3e). We extracted 1,000 bp upstream promoter sequences of these genes from the TAIR10 database and performed *de novo* motif discovery using STREME. This analysis identified several significantly enriched motifs (Figure S2a). Among these, we noted that motifs such as ERSE-I, MYB/MADS binding motifs, an ACTACCC motif, a NAC binding motif (CCAGAAACA), and a WRKY binding motif (AATGTCAAC) were particularly enriched in the promoters of genes belonging to cluster 6, which exhibited pronounced differences in expression between *bzip28/60* and *coffin1* under TM treatment (Figure 3e, f). Given the known roles of NAC and WRKY TFs in stress and defense responses^25,27,29,36,37^, we focused on these two TF families for further investigation. To narrow down candidate TFs, we cross-referenced the promoters of the UPR genes in cluster 6 with a publicly available DNA affinity purification sequencing (DAP-seq) database^38^. This analysis identified six candidate TFs—one NAC (NAC2) and five WRKYs (WRKY8, WRKY15, WRKY22, WRKY28, WRKY45)—that are known to bind to at least one of the promoters of the upregulated UPR genes (Figure S2b). The enrichment of NAC and WRKY binding motifs in the promoters of UPR genes, particularly in cluster 6, suggests that these TF families play a role in modulating UPR gene expression during ER stress, potentially downstream of MORG1.

### WRKY8 negatively regulates UPR gene expression

To determine which of the candidate TFs identified in the promoter analysis (Figure S2b) suppress UPR gene expression, we cloned the coding sequences of six of these TFs into expression vectors driven by the Cauliflower Mosaic Virus (CaMV) 35S promoter. Using dual-luciferase assays in *Nicotiana tabacum* leaves^14^, we tested these TFs for their regulatory effects on a 1 kb *BiP3* promoter fused to firefly luciferase (FLUC) as the reporter, with *35S::Renilla luciferase* (RLUC) serving as the internal control. Co-expression of constitutively active forms of bZIP28 (truncated bZIP28 lacking the transmembrane domain)^14^ and spliced bZIP60 (sbZIP60)^14^ with the vector control robustly activated the *BiP3* promoter, confirming the functionality of the reporter system (Figure 4a). Among the tested TFs, only WRKY8 significantly reduced *BiP3* promoter activity when co-expressed with active bZIP28 and bZIP60, indicating its role as a repressor of UPR gene expression (Figure 4a, Figure S3). The other five candidate TFs did not significantly reduce *BiP3* promoter activity under these conditions (Figure S3). Interestingly, the expression of *MORG1* also decreased *BiP3* promoter activity. However, co-expression of *MORG1* and *WRKY8* did not further reduce activity compared to *WRKY8* or *MORG1* alone, suggesting that they likely function within the same regulatory pathway, with MORG1 potentially modulating WRKY8 activity (Figure 4a).

**Figure 4.**
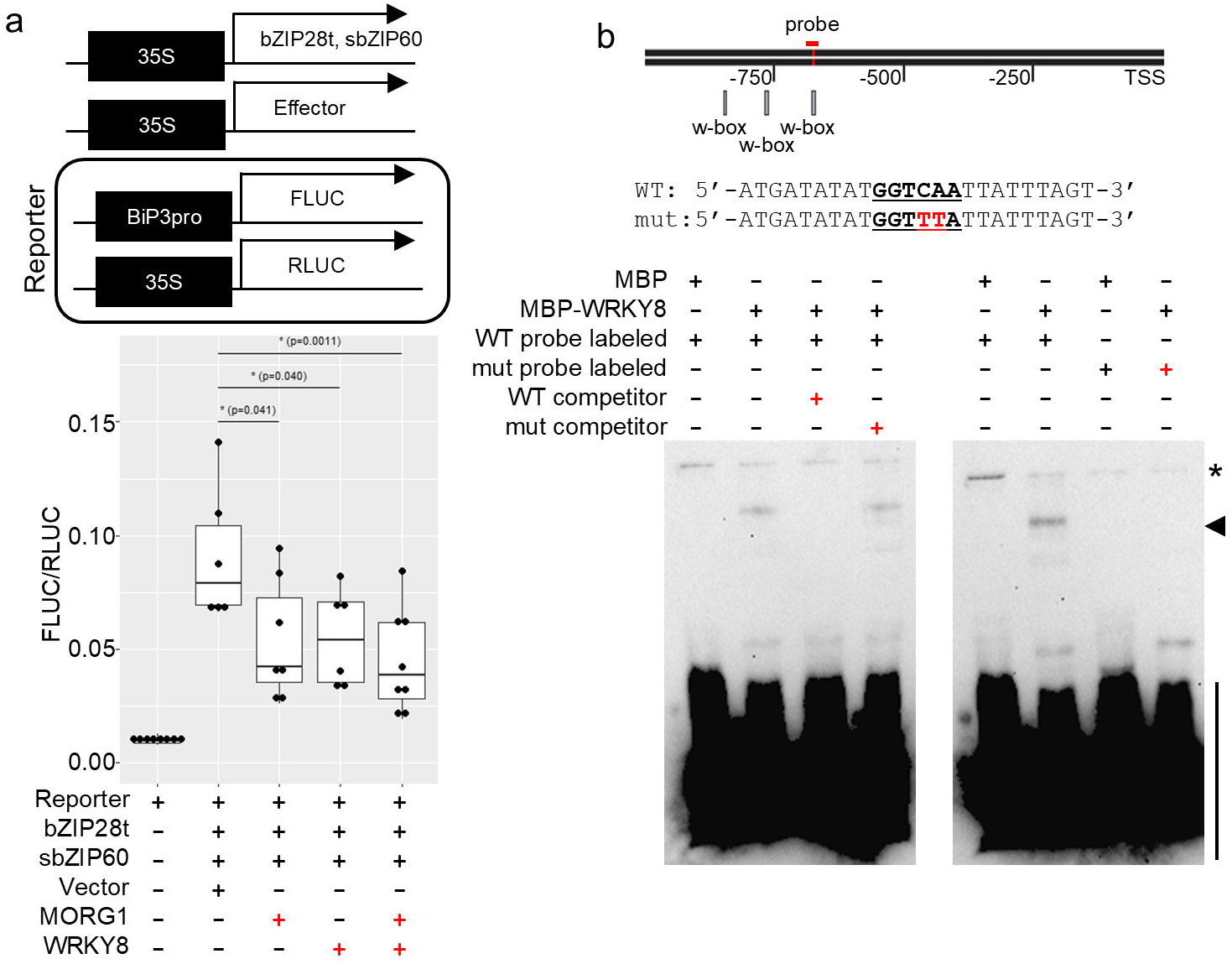
WRKY8 represses *BiP3* promoter activity. **a.** Dual-luciferase assay to assess the effect of MORG1 and WRKY8 on BiP3 promoter activity. (Top) Schematic diagram illustrating the dual-luciferase assay procedure. (Bottom) BiP3 promoter activity in the presence of truncated bZIP28, spliced bZIP60, and either a vector control or an effector (MORG1 or WRKY8). *Nicotiana tabacum* leaves were agroinfiltrated with the indicated constructs. Plus signs indicate addition of the corresponding Agrobacterium cell culture. Data were normalized to Renilla. Median ± IQR.; *n* = 6-8 biological replicates. Statistical significance was determined by Student’s *t*-test. **b**. Electrophoretic mobility shift assay (EMSA) demonstrating WRKY8 binding to the BiP3 promoter. (Top) Schematic diagram of the BiP3 promoter region, indicating the location of the WRKY binding motif used for probe design. Sequences of the wild-type (WT) and mutated probes are shown. (Bottom) EMSA results using biotin-labeled probes. Recombinant WRKY8 protein was incubated with the labeled probes, and the resulting complexes were resolved by electrophoresis. Plus signs indicate addition of the corresponding component. Competitor DNA was added at 200x concentration of the labeled probe. Arrowheads indicate DNA-protein complexes. Asterisks indicate samples retained in the well.

### WRKY8 directly binds to the *BiP3* promoter

To investigate whether WRKY8 directly interacts with UPR gene promoters, we performed electrophoretic mobility shift assays (EMSA)^39,40^ using recombinant WRKY8 protein and biotin-labeled DNA probes containing W-box elements from the *BiP3* promoter. WRKY8 specifically bound the *BiP3* promoter probe, forming a distinct DNA–protein complex (Figure 4b). This binding was effectively competed by an excess of unlabeled probe, but not by a mutated probe lacking the W-box sequence (Figure 4b). These results confirm that WRKY8 directly binds to the *BiP3* promoter through the W-box motif, establishing its role as a transcriptional regulator of UPR genes. Furthermore, using Plant PAN 4.0^41^, we analyzed the promoter sequences of six UPR genes (*BI1*, *BiP3*, *ERdj3A*, *ERdj3B*, *PDIL1*-2, *PDIL2*-2) and found that all of them contain at least one putative W-box element within 1 kb upstream of the transcription start site (Figure S4a). Analysis of a publicly available ChIP-seq database (PCBase2)^41^ also indicated that WRKY TFs bind to the promoter regions of these UPR genes *in vivo* (Figure S4b). While this ChIP-seq dataset was generated under MAMP-triggered immunity conditions and not under ER stress conditions, and the binding of WRKY8 was not specifically analyzed^42^, together, these findings suggest that WRKY TFs, potentially including WRKY8, may directly regulate not only *BiP3* but also other UPR genes by binding to their W-box-containing promoters.

### MORG1 modulates MPK6-mediated phosphorylation of WRKY8

MORG1 does not have kinase activity but functions as a MAP kinase scaffold protein^21,23^. Therefore, we hypothesized that it influenced the activation of MAP kinases, including MPK3 and MPK6, which are known to play crucial roles in plant stress signaling. To test this hypothesis, we first examined the phosphorylation status of MPK3 and MPK6 in WT, *bzip28/60*, and *coffin1* seedlings under TM treatment using phospho-specific antibodies.

A basal level of MPK3/6 phosphorylation was observed in all genotypes under mock (DMSO) conditions, likely due to the seedling handling in the assays (Figure 5a). However, upon TM treatment but not in the control, MPK3 and MPK6 phosphorylation increased over time, peaking around 2 hours. Notably, the TM-induced phosphorylation of MPK3/6 was significantly reduced in *coffin1* mutants compared to WT and *bzip28/60*, suggesting that MORG1 is required for proper activation of MPK3 and MPK6 (Figure 5a).

**Figure 5.**
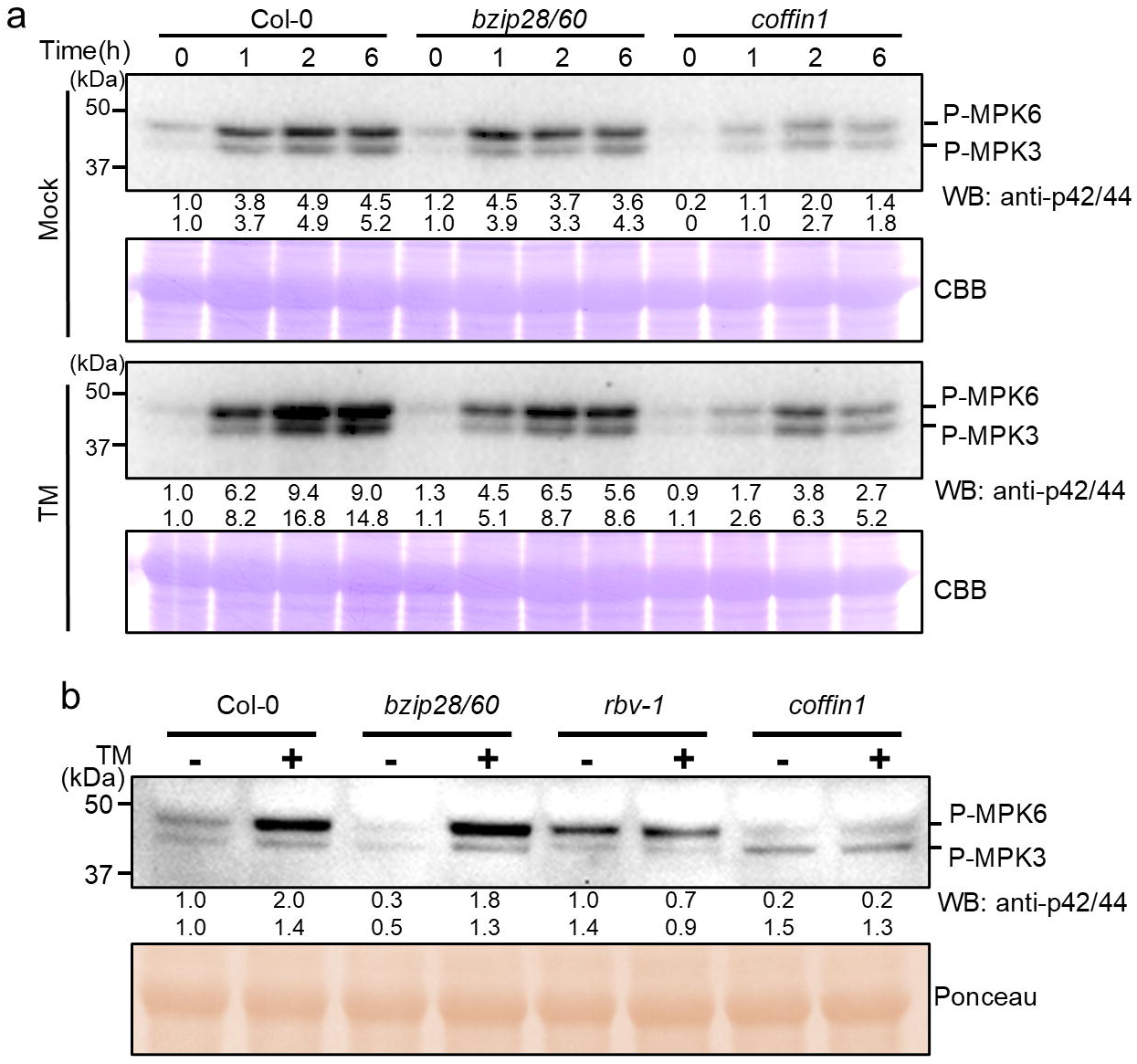
Disruption of MORG1 attenuates TM-induced MPK3/6 phosphorylation. **a.** Western blot analysis of MPK3/6 phosphorylation in response to ER stress over time. Col-0, *bzip28/60*, and *coffin1* seedlings were treated with DMSO (Mock) or 500 ng/ml TM for the indicated times (0, 1, 2, and 6 hours). The upper band corresponds to phosphorylated MPK6 (p-MPK6), and the lower band corresponds to phosphorylated MPK3 (p-MPK3). Numbers below the blot indicate the relative intensity of each band, quantified using ImageJ and normalized first to the loading control and then to the first lane. Coomassie Brilliant Blue (CBB) staining of the blot is shown below as a loading control. **b.** Western blot analysis of MPK3/6 phosphorylation in response to ER stress. Col-0, *bzip28/60*, *rbv-1*, and *coffin1* seedlings were treated with DMSO (Mock) or 500 ng/ml TM for 2 hours. The upper band corresponds to phosphorylated MPK6 (p-MPK6), and the lower band corresponds to phosphorylated MPK3 (p-MPK3). Numbers below the blot indicate the relative intensity of each band, quantified using ImageJ and normalized first to the loading control and then to the first lane. Ponceau S staining of the blot is shown below as a loading control.

To further confirm the role of MORG1 in regulating MPK3/6 activation, we compared the phosphorylation levels of these proteins in WT, *bzip28/60*, *rbv-1*, and *coffin1* seedlings after 2 hours of TM treatment. Both *coffin1* and *rbv-1* mutants exhibited significantly reduced MPK3/6 phosphorylation compared to WT and *bzip28/60* under TM treatment (Figure 5b). Interestingly, while the ratio of phosphorylated MPK6 (p-MPK6) to phosphorylated MPK3 (p-MPK3) was higher in WT, *bzip28/60*, and *rbv-1*, the ratio was reversed in *coffin1*, with p-MPK3 levels exceeding p-MPK6 levels. Although the exact mechanism underlying this altered ratio remains unclear, it may reflect a differential impact of the *coffin1* mutation on MPK3 and MPK6 activation or stability (Figure 5b).

Given that MPK3/6 phosphorylate downstream targets, including TFs, we hypothesized that MORG1 influences WRKY8 phosphorylation by modulating MPK3/6 activity. To directly test whether MORG1 impacts WRKY8 phosphorylation, we conducted *in vitro* kinase assays. Recombinant WRKY8 protein was incubated with MPK6 in the presence or absence of MORG1. Phosphorylation was analyzed by Phos-tag SDS-PAGE, which allows for the separation of differentially phosphorylated forms of a protein^43^. MPK6 alone was able to phosphorylate WRKY8 (Figure 6g). However, the addition of MORG1 led to the appearance of an additional band, indicating that MORG1 enhances the phosphorylation of WRKY8 by MPK6, possibly targeting additional sites or altering stoichiometry (Figure 6g). These findings suggest that MORG1 functions as a scaffold, facilitating the recruitment of MPK6 and WRKY8 to enable efficient and potentially distinct phosphorylation patterns.

**Figure 6.**
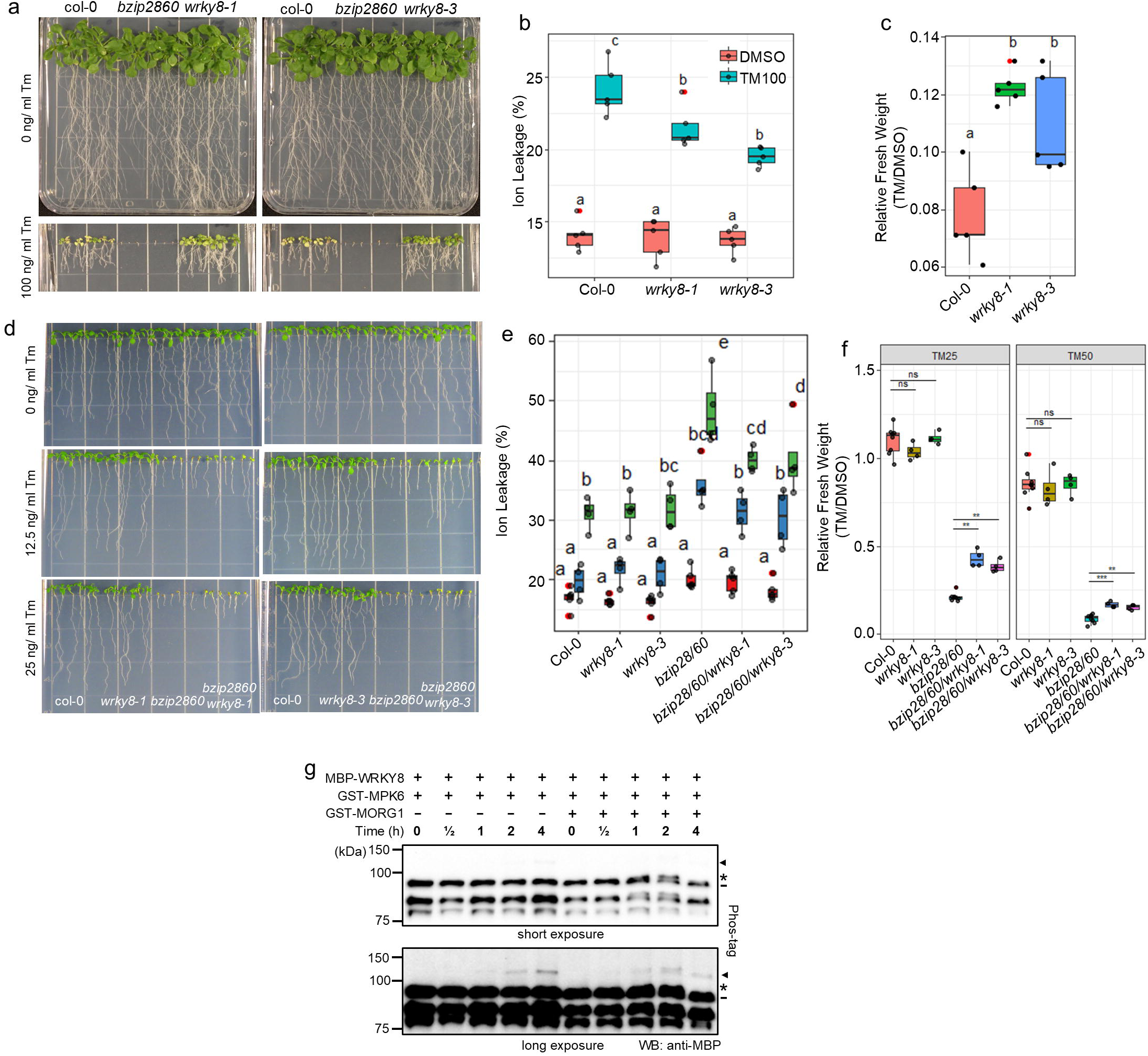
WRKY8 is phosphorylated by MAP kinase cascade under ER stress and is a pro-death factor for ER stress. **a.** Phenotypic analysis of *wrky8* mutants under ER stress. Col-0, *bzip28/60*, *wrky8-1*, and *wrky8-3* seedlings were grown on 1/2 LS media supplemented with DMSO (Mock) or 100 ng/ml TM. Representative images of seedlings are shown. **b**. Ion leakage of col-0, *wrky8-1*, and *wrky8-3* seedlings under ER stress conditions. Seedlings were grown as described in **a**. Median ± IQR.; *n* = 4 replicates (seedlings pooled per replicate). Statistical significance was determined by one-way ANOVA with Tukey’s post-hoc test. **c**. Relative fresh weight of col-0, *wrky8-1*, and *wrky8-3* seedlings under ER stress conditions.. Median ± IQR.; *n* = 5 replicates (seedlings pooled per replicate). Statistical significance was determined by one-way ANOVA with Tukey’s post-hoc test. **d**. Phenotypic analysis of *bzip28/60 x wrky8-1* and *bzip28/60 x wrky8-3* mutants under ER stress. Seedlings were grown on 1/2 LS media supplemented with DMSO (Mock), 12.5 ng/ml, or 25 ng/ml TM. Representative images of seedlings are shown. **e**. Ion leakage of seedlings shown in **d** under ER stress conditions. Median ± IQR.; *n* = 4 replicates (10 seedlings pooled per replicate). Statistical significance was determined by one-way ANOVA with Tukey’s post-hoc test. **f**. Relative fresh weight of seedlings shown in **d** under ER stress conditions. Median ± IQR.; *n* = 5 replicates (seedlings pooled per replicate). Statistical significance was determined by one-way ANOVA with Tukey’s post-hoc test. **g**. *In vitro* kinase assay demonstrating WRKY8 phosphorylation by MPK6. MBP-WRKY8 was incubated with GST-MPK6 in the presence or absence of GST-MORG1 for the indicated times (0, 30 min, 1, 2, and 4 hours). Reaction products were analyzed by Western blotting using an anti-MBP antibody. Arrowheads indicate phosphorylated WRKY8 by MPK6 alone bands. The asterisk indicates differently phosphorylated WRKY8. Dash mark indicates MBP-WRKY8.

### *Wrky8* mutants exhibit enhanced ER stress tolerance and UPR gene expression

Our results thus far point to the role of WRKY8 in UPR gene expression and likely cell fate in unresolved ER stress. Therefore, to validate the role of WRKY8 in ER stress responses, we analyzed two established independent T-DNA insertion alleles, *wrky8-1* and *wrky8-3*, which disrupt WRKY8 expression^30^. Under chronic TM-induced ER stress, both *wrky8* alleles displayed higher fresh weight, and reduced ion leakage compared to WT plants (Figure 6a–c). These phenotypes were similar to those observed in *coffin1* and *rbv-1* mutants (Figure 2k-i), suggesting that loss of WRKY8 enhances ER stress tolerance. Next, to explore a functional relationship of WRKY8 with the bZIP28/60 pathway, we generated a *bzip28/60/wrky8* triple mutant. Under TM treatment, the triple mutant exhibited increased growth compared to *bzip28/60* (Figure 6d–f). These results support that WRKY8 functions downstream of bZIP28 and bZIP60 to repress the cytoprotective role of these TFs in ER stress.

## Discussion

In this study, we identify the MAP kinase scaffold protein MORG1 and the TF WRKY8 as novel negative regulators of the UPR and ER stress tolerance operating in a novel functional pathway controlling UPR gene expression (Figure 7). Specifically, through an EMS suppressor screen in the hypersensitive *bzip28/60* double mutant background, we isolated the *coffin1* mutant, which exhibited enhanced tolerance to chronic ER stress (Figure 1). Genetic and molecular analyses revealed that the *coffin1* phenotype arises from a mutation in MORG1 (Figure 2), implicating this MAP kinase scaffold protein in regulating life-or-death decisions led by the UPR. Our findings demonstrate that MORG1 negatively regulates ER stress responses by modulating the phosphorylation of the transcription factor WRKY8 via MPK6. We show that MORG1 is required for proper MPK3/6 activation under ER stress, as evidenced by the reduced phosphorylation of these kinases in the *rbv-1* and *coffin1* mutant (Figure 5), suggesting that MORG1 facilitates kinase activation by potentially recruiting upstream MKKs to MPK3/6. In the absence of MORG1, compromised MPK3/6 activation leads to altered downstream signaling, with significant implications for stress response regulation.

**Figure 7.**
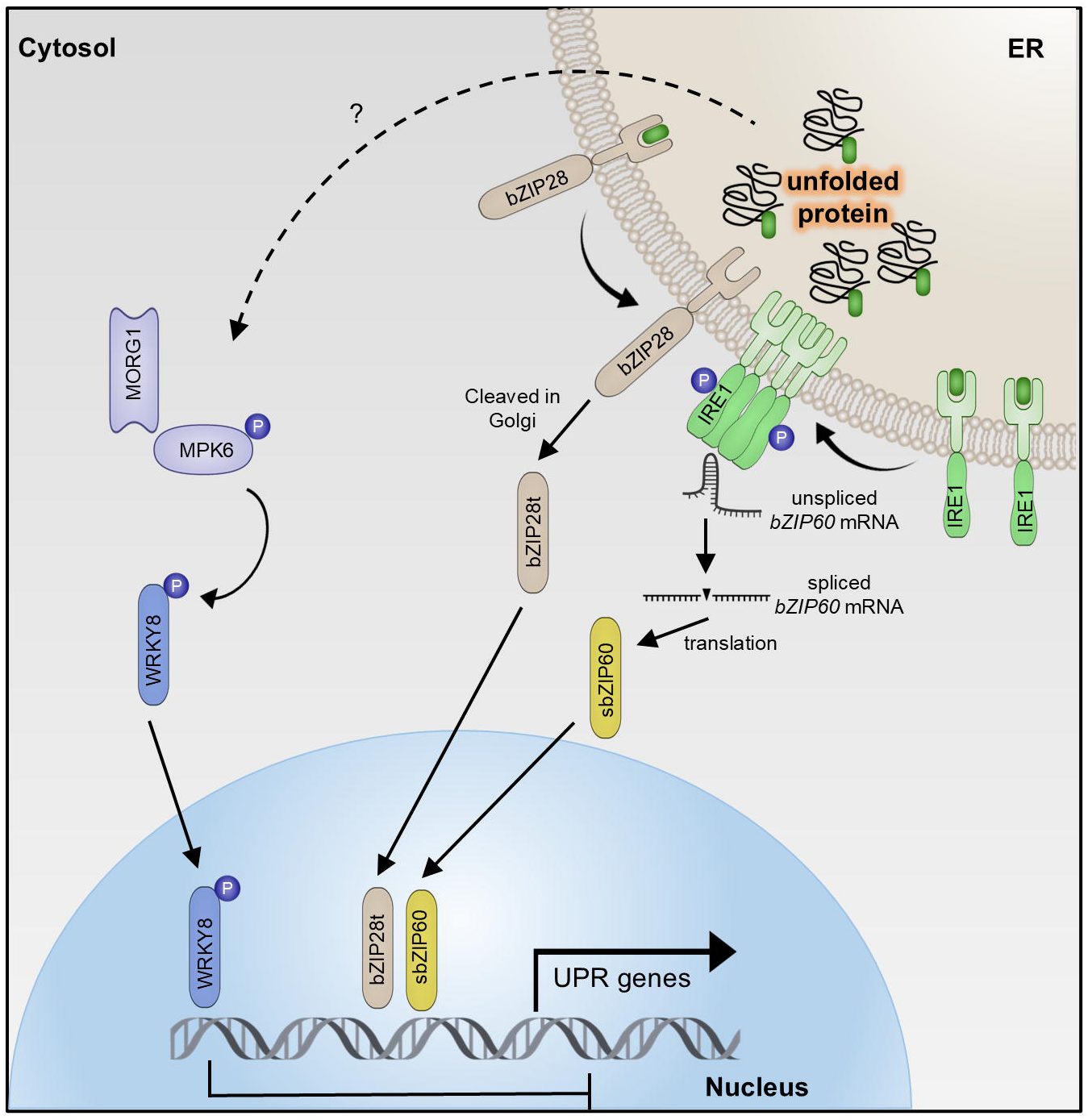
Proposed model for the role of MORG1 in modulating the UPR and cell death under unresolved ER stress. Upon ER stress, misfolded proteins accumulate in the ER lumen and are sensed by IRE1, which then splices *bZIP60* mRNA to produce the active, spliced form (sbZIP60). In parallel, bZIP28 is transported to the Golgi apparatus, where it is cleaved to release its cytosolic domain (bZIP28t). Both sbZIP60 and tbZIP28 translocate to the nucleus and activate the transcription of UPR target genes, promoting ER stress adaptation and cell survival. This study reveals a novel regulatory pathway involving the MAP kinase scaffold protein MORG1, which modulates the UPR through a previously unknown mechanism. Under ER stress, MORG1 facilitates the activation of MPK6, which in turn phosphorylates the transcription factor WRKY8. Phosphorylated WRKY8 exhibits enhanced DNA binding affinity to the promoters of UPR genes, including BiP3, and represses their expression. In the *morg1* mutant, reduced MPK6 activation leads to decreased WRKY8 phosphorylation, thereby relieving the repression of UPR genes and promoting cell survival under unresolved ER stress. The upstream signal that triggers the MORG1-MPK6-WRKY8 pathway remains to be determined. This MORG1-MPK6-WRKY8 pathway acts in parallel to the canonical IRE1-bZIP60 and bZIP28 pathways, providing a novel mechanism for fine-tuning the UPR and influencing cell fate decisions under ER stress.

Our discoveries add ER stress responses to the expanding functional repertoire of MORG1, which was previously known for its roles in facilitating ERK signaling in mammals^23^, regulating mTORC1 activity under nutrient stress^22^, and mediating defense responses through the GhMKK6-GhMPK4 cascade in cotton^21^. In Arabidopsis, MORG1 was implicated in microRNA biogenesis by promoting ARGONAUTE1 loading^35^. Our study also identifies WRKY8 as a key negative regulator of UPR gene expression, contrasting with the activating roles of bZIP28 and bZIP60 in UPR gene regulation. We show that WRKY8 directly binds to W-box motifs in the promoters of *BiP3* (Figure 4b) and potentially other UPR genes, including *BI1*, *ERdj3A*, *ERdj3B*, *PDIL1*-2, and *PDIL2-2* (Figure S4a, S4b).

Our findings that WRKY8 has a critical role in ER stress resonate with previous studies on other WRKY family members. For instance, WRKY7, -11, and -17 were shown to suppress the bZIP28-dependent UPR pathway during PAMP-triggered immunity^27^, suggesting a broader role for WRKY TFs in modulating ER stress responses across different contexts. While our study focuses on WRKY8 in ER stress, further research could explore whether these WRKYs act on similar UPR gene targets or through distinct yet potentially interconnected pathways. The negative regulation of UPR genes exerted by WRKY8 likely attenuates ER stress responses once ER homeostasis is restored or prevents excessive activation that could result in programmed cell death. This fine-tuning mechanism, potentially involving multiple WRKY TFs, ensures a balance between alleviating ER stress and avoiding the metabolic costs of sustained UPR activation. Furthermore, the specific involvement of WRKY8 in ER stress adds another layer to its already established roles in biotic stress responses, including pathogen infection and wounding, as well as abiotic stress responses like salinity tolerance. These considerations suggest that WRKY8 may be a crucial node integrating various stress signals to modulate appropriate downstream responses.

Our *in vitro* kinase assays and genetic analyses support a model in which MORG1 facilitates MPK6-mediated phosphorylation of WRKY8 (Figures 5, 6g). In the *morg1* mutants (*rbv-1, coffin1*), reduced WRKY8 phosphorylation diminishes its repressor function, leading to partial restoration of UPR gene expression in the absence of bZIP28 and bZIP60 (Figure 3e-g). This mechanism provides an alternative pathway for modulating UPR genes independent of the canonical bZIP28 and bZIP60-mediated UPR pathway. The partial restoration of UPR gene expression in *coffin1* and *bzip28/60/wrky8* mutants suggests that alternative pathways can compensate for the loss of bZIP28 and bZIP60 under specific conditions. This also suggests that the MORG1-MPK6-WRKY8 pathway may act in parallel with or downstream of the canonical UPR pathways to fine-tune the ER stress response.

Our study opens the door to additional significant research. First, identifying the upstream signals that activate the MORG1–MPK6–WRKY8 pathway during ER stress will be crucial for understanding how this mechanism integrates into the broader stress response network. Potential candidates include ER stress sensors or cross-talk with other signaling pathways. Second, the role of other WRKY TFs in UPR regulation warrants investigation. Functional redundancy or interplay among WRKY family members could reveal additional layers of complexity. Lastly, examining the conservation of the regulatory mechanism identified in the work among multicellular eukaryotes could pave the way for translational applications to improve organismal resilience under stress conditions.

In conclusion, our study establishes MORG1 as a critical modulator of ER stress responses in Arabidopsis, functioning through the regulation of WRKY8 phosphorylation by MPK6. This novel pathway exemplifies how scaffold proteins, MAP kinases, and TFs interact to shape plant stress signaling networks and provides a foundation for developing strategies to enhance stress tolerance in crops facing increasingly challenging environmental conditions.

## MATERIALS AND METHODS

### Plant materials and growth conditions

All seeds were sown on ½ Linsmaier Skoog (LS) Basal Medium (Caisson Labs) supplemented with 1% Sucrose, 1.2% Agar, and DMSO or designated concentration of Tunicamycin. After stratification in the dark at 4℃ for 2 days, plates were transferred to a growth chamber with 80 µmol m^-2^s^-1^ under 16 h light / 8 h dark at 22 ℃.

### Plasmid construction

For *coffin1* complementation, the MORG1 construct was generated by amplifying a 1 kb MORG1 promoter along with the MORG1 coding sequence (CDS). The StayGold fluorescent protein^44^ was fused to the C-terminal end of the MORG1 CDS. This fragment was first cloned into the pDONR207 vector using Gateway™ BP Clonase™ II Enzyme mix (Thermo Fisher Scientific) and subsequently transferred into the pGWB1 destination vector through an Gateway™ LR Clonase™ II Enzyme mix (Thermo Fisher Scientific).

For the dual luciferase assay^16,45^, the coding sequences of MORG1, MPK3, MPK6, and MORG1 target candidate genes were amplified from cDNA derived from Col-0 plants. These sequences were cloned into the pGreenII 62SK vector^46^ for expression.

For the production of recombinant proteins, the MORG1 and MPK6 coding sequences were cloned into the pDONR207 entry vector using Gateway™ BP Clonase™ II Enzyme mix (Thermo Fisher Scientific) and then transferred to the pDEST15 vector (Thermo Fisher Scientific) using Gateway™ LR Clonase™ II Enzyme mix. The MBP-WRKY8 construct was generated by cloning the corresponding coding sequence into the pMAL-c5x vector (New England Biolabs) using restriction enzyme digestion and ligation.

### Recombinant protein expression and purification

The GST-MORG1 and GST-MPK6 proteins were expressed and purified according to the manufacturer’s instructions (GoldBio) with minor modification. E. coli (BL21) cells were grown in a shaking incubator at 18°C for 16 hours after induction with 0.3 mM isopropyl-β-d-thiogalactoside (IPTG) at an optical density at 600 nm (OD600) of 0.6–0.8. The cells were harvested by centrifugation at 6000 rpm for 10 minutes at 4°C. The resulting pellet was resuspended in lysis buffer (50 mM Tris-HCl [pH 8.0], 150 mM NaCl, 1 mM EDTA, 0.1% Triton X-100, and 2 mM phenylmethylsulfonyl fluoride [PMSF]) and lysed by ultrasonication. GST-tagged fusion proteins were purified using Glutathione Agarose Resin (GoldBio) and eluted with buffer containing 10 mM reduced glutathione. The purified proteins were used in both the EMSA and in vitro kinase assays.

The MBP-WRKY8 protein was expressed and purified following the manufacturer’s protocol (New England BioLabs) with minor modification. E. coli (BL21) cells were cultured in a shaking incubator at 18°C for 16 hours after induction with 0.3 mM IPTG at an OD600 of 0.6–0.8. Harvested cells were resuspended in lysis buffer (20 mM Tris-HCl [pH 7.4], 200 mM NaCl, 1 mM EDTA, and 2 mM PMSF) and disrupted by ultrasonication. MBP-tagged fusion proteins were purified using amylose resin (New England BioLabs) and eluted with buffer containing 10 mM maltose. The purified proteins were utilized in EMSA and in vitro kinase assays.

### EMS mutant generation

The procedure for the EMS mutagenesis followed a previous paper^47^. In brief, 20,000 *bzip28/60* seeds were treated with 25 volumes of 0.3% (v/v) EMS (Sigma-Aldrich) in a 50-ml conical tube for 15 h with rotation. The EMS solution was discarded using a pipette. After eight washes with distilled water, seeds were soaked in distilled water for 1 h to allow the EMS to diffuse out from the seeds. M_1_ seeds were sown on soil and grown under the growth conditions described above. M_2_ seeds were collected from individual M_1_ plants (>500 lines) and >200,000 M_2_ seeds were screened for chronic ER stress resistance.

### Bulked Segregant Analysis

To perform BSA, we created an F_2_ population (BC_1_F_2_) derived from a backcross between *coffin1* and *bzip28/60*. Genomic DNA from pools of ≥ 300 *coffin1* BC_1_F_2_ whole seedlings with or without the suppressor phenotype and ≥ 300 M_4_ *coffin1* whole seedlings was extracted using NucleoSpin Plant Midi kit (MACHEREY-NAGEL, Düren, Germany) according to the manufacturer’s instruction. The final purified genomic DNA was quantified using both Qubit dsDNA HS (Thermo Fisher Scientific, Carlsbad, CA) and Agilent 4200 TapeStation High Sensitivity DNA 1000 assays (Agilent Technologies, Santa Clara, CA), and then subjected to library construction using the TruSeq Nano DNA Library Preparation Kit (Illumina, San Diego, CA). The libraries were sequenced in pair-end mode on the Illumina NovaSeq 6000 platform (150-nt) at the Research Technology Support Facility (RTSF) Genomics Core at Michigan State University. The quality of raw reads was evaluated using FastQC (version 0.11.5). Reads were cleaned for quality and adapters with Cutadapt^48^ (version 1.8.1) using a minimum base quality of 20 retaining reads with a minimum length of 30 nucleotides after trimming. Quality-filtered reads were aligned to the Col-0 reference genome (TAIR10) using Bowtie^49^ (version 2.2.3). The concordantly mapped reads with ≥ 10 of the mapping score were extracted using a custom script and retained for further analysis. Duplicated reads were subsequently removed using Samtools^50^ (version 1.8). Variant calling was performed using Samtools63 (version 1.8) and recorded in the VCF using bcftools (version 1.9.64). The resulting vcf files were converted into the format required for SHOREmap analysis^51^ (version 3.6) using “SHOREmap convert”. The consensus information for candidate markers was extracted using “SHOREmap extract”. Allele frequency of the candidate markers that had ≥ 25 marker score and ≥ 10 of coverage in the population was analyzed using “SHOREmap backcross” with a background correction (--bg-freq 0.4). The filtered candidate markers were annotated to TAIR10 using “SHOREmap annotate”. The genome coverage information of our BSA is provided in Supplementary Table 1. A full list of markers obtained through SHOREmap is provided in Supplementary Data 1.

### Reverse transcription-quantitative PCR (RT-qPCR)

Total RNA was extracted from 5-day-old seedlings treated with 0.5 µg/ml Tm or DMSO for 6 h using the NucleoSpin RNA Plant kit (Macherey-Nagel) according to the manufacturer’s instructions. cDNA was synthesized from 500 ng of RNA using the iScript™ cDNA Synthesis Kit (Bio-rad). Fast SYBR Green Master Mix (Applied Biosystems) was used in the presence of gene-specific primers and template cDNAs in an ABI7500 (Applied Biosystems). The list of primers used in qRT–PCR with gene accessions is provided in Supplementary Table 1.

### Western blot analysis

For the analysis of MPK3/6 phosphorylation, total proteins were extracted from samples harvested under the conditions described in the main text. Seedlings were frozen in liquid nitrogen and finely ground. The samples were then resuspended in lysis buffer (1% (w/v)SDS, 10 mM Tris (pH8.0), 1 mM EDTA) and centrifuged at 13,000 rpm for 5 min. The supernatant was collected, and total protein concentration was determined using the DC protein assay (Bio-rad). 50 µg of total protein was loaded per well and resolved by 10% SDS-PAGE. The primary antibody, rabbit anti-42/44 antibody (Cell Signaling Technology), was used at a 1:1000 dilution for Western blot analysis.

For in-gel kinase assay samples, proteins were resolved using 8% phos-tag (Fujifilm Wako) SDS-PAGE^52^. For Western blot analysis, the mouse anti-MBP antibody (New England Biolabs) was used at a 1:4000 dilution.

### RNA seq data analysis

RNA-seq libraries were constructed using the Illumina TruSeq Stranded mRNA Library (Illumina, San Diego, CA, USA) and sequenced in paired-end mode on the Illumina NovaSeq 6000 platform (150-nt) at the RTSF Genomics Core at Michigan State University. The quality of raw reads was assessed using FastQC (version 0.11.5). Reads were cleaned for quality, and adapters with Cutadapt^48^ (version 1.8.1) were used, with a minimum base quality of 20 retaining reads and a minimum length of 30 nucleotides after trimming. Quality-filtered reads were aligned to the Col-0 reference genome (TAIR10) using Bowtie^49^ (version 2.2.4) and TopHat^53^ (version 2.0.14) with a 10-bp minimum intron length and 15,000-bp maximum intron length. Fragments per kilobase exon model per million mapped reads (FPKM) were calculated using TAIR10 gene model annotation with Cufflinks^54^ (version 1.3.0). Per-gene read counts were measured using HTSeq^55^ (version 0.6.1p1) in the union mode with a minimum mapping quality of 20 with stranded=reverse counting. Differential gene expression analysis was performed in each sample relative to the mock control using DESeq2^56^ (version 1.36.1) within R (version 4.1.3). Genes of which the total count across treatments and replicates in each genotype is < 100 were not included in the analysis. All genes analyzed were visualized for each genotype in volcano plots using the R package EnhancedVolcano (version 1.18). DEGs were obtained based on adjusted P-value < 0.05 and absolute Log2FC > 1. GO enrichment analysis was performed using agriGO^57^ (version 2.0) (http://systemsbiology.cau.edu.cn/agriGOv2/) with a false-discovery rate adjusted P < 0.05 (hypergeometric test with Bonferroni correction) as a cutoff. Biological process GO categories were analyzed and visualized in the heatmap using the R package ComplexHeatmap^58^ (version 2.14.0). K-means clustering analysis of the 1,215 DEGs was performed with Log2FC outputs generated from DESeq2 using R package factoextra (version 1.0.7).

### Cistrome analysis

The 1-kb upstream sequences of the transcription start site of Cluster 1 (the sequences of 247 DEGs available out of 250 DEGs), Cluster 2 (102 DEGs), Cluster 3 (185 DEGs), Cluster 4 (336 DEGs), Cluster 5 (165 DEGs), Cluster 6 (34 DEGs), Cluster 7 (98 DEGs), and Cluster 8 (38 DEGs) were obtained from the BioMart tool in the Phytozome database (version 13). The promoter sequences of the identical number of randomly selected genes for each cluster were also obtained for control sets. De novo motif discovery in each cluster was performed on the 1-kb promoters using STREME^59^ with a parameter of “-objfun de --dna -- nmotifs 15 --minw 7 --maxw 20” with the control set of random genes. The resulting data was visualized in the heatmap using the R package ComplexHeatmap^58^ (version 2.14.0).

### Dual Luciferase assay

The pGreenII 0800-LUC-BiP3 promoter vector and truncated pGreenII 62-SK bZIP28 (bZIP28t), spliced bZIP60 (sbZIP60) were acquired from a previous study^14^. *Nicotiana tabacum* leaves were infiltrated with *Agrobacterium* suspensions carrying the indicated constructs. *Agrobacterium* cell cultures were prepared at an OD600 of over 1.0 and diluted to 0.1 in agroinfiltration buffer (10 mM MgCl2, 10 mM MES (pH 5.6), 200 μM acetosyringone). The cultures were then incubated in the dark at room temperature for 4 hours before infiltration. Three days after infiltration, leaf discs were collected, and luciferase activity was measured using the Dual-Luciferase Reporter Assay System (Promega). Firefly luciferase (FLUC) activity was normalized to Renilla luciferase (RLUC) activity.

### Electrophoretic mobility shift assay (EMSA)

EMSA was performed using the LightShift Chemiluminescent EMSA Kit (Thermo Scientific) following the manufacturer’s instructions. Recombinant MBP and MBP-WRKY8 proteins were used for binding assays. Biotin-labeled DNA probes (Eurofins Genomics) were synthesized for the wild-type (WT) and mutant (mut) sequences of the target DNA. Binding reactions were carried out in 20 µL volumes containing 1X binding buffer (100 mM Tris, 500 mM KCl, 10 mM DTT; pH 7.5), 50 ng/μL poly(dI•dC), 5 mM MgCl₂, 2.5% glycerol, 20 fmol of biotin-labeled probe, and the specified protein concentration. For competition assays, a 200-fold molar excess of unlabeled WT or mutant competitor DNA was included.

Binding reactions were incubated at room temperature for 30 minutes and then mixed with 5X loading buffer before being electrophoresed on a 6% TBE polyacrylamide gel in 0.5X TBE buffer. Following electrophoresis, DNA-protein complexes were transferred onto a positively charged nylon membrane (GE healthcare) and crosslinked using UV light (254 nm, 120,000μ joules). Detection of biotin-labeled DNA was performed using the Streptavidin-Horseradish Peroxidase Conjugate included in the kit and chemiluminescent substrate according to the manufacturer’s protocol. The chemiluminescence signal was captured using a ChemiDoc MP imager (Bio-Rad).

### *In vitro* kinase assay

*In vitro* kinase assays were performed using recombinant MBP-WRKY8, GST-MPK6, and GST-MORG1 proteins. The reactions were conducted in a final volume of 100 µL containing kinase reaction buffer^43^ (25 mM Tris-HCl [pH 7.5], 12 mM MgCl₂, 1 mM DTT, and 1 mM ATP). Protein components were added as follows: MBP-WRKY8 (4 µg), GST-MPK6 (2 µg), and GST-MORG1 (2 µg). The reaction mixtures were incubated at 30°C for the specified duration. Following incubation, the reactions were stopped by adding 6X SDS sample buffer and heating at 95°C for 5 minutes. Phosphorylated proteins were separated on a Phos-tag SDS-PAGE gel. After electrophoresis, the proteins were transferred to a PVDF membrane and analyzed by western blotting.

### Electrolyte leakage measurement

Cell death was determined by quantification of percent electrolyte leakage as described previously^60,61^ with minor modifications. Ten seedlings per sample were rinsed thoroughly with deionized water and placed in 2 mL of Milli-Q water. The tubes were incubated at room temperature for 4 hours with gentle shaking. After incubation, the conductivity of the solution was measured using a conductivity meter to determine the electrolyte leakage. Subsequently, the tubes containing the seedlings and incubation solution were autoclaved to release the total electrolytes. After cooling to room temperature, the conductivity of the solution was measured again. Electrolyte leakage was calculated as the percentage of initial conductivity relative to total conductivity.

## Data availability

All data supporting the findings of this study are available within this paper and its Supplementary Materials files. The raw data of WGS and RNA-seq have been deposited to the National Center for Biotechnological Information Sequence Read Archive and are accessible via BioProject accession codes PRJNA1196415 and PRJNA1196845, respectively. The Col-0 reference genome (TAIR10) was used for sequence analyses. The DAP-seq dataset was downloaded from the Plant Cistrome database (http://neomorph.salk.edu/dev/pages/shhuang/dap_web/pages/index.php).

## Code availability

The scripts used in this study are available in GitHub (https://github.com/DaeKwan-Ko/coffin1).

## Acknowledgments

This study was supported primarily by the National Institutes of Health (R35GM136637) with contributing support from by the Great Lakes Bioenergy Research Center, U.S. Department of Energy, Office of Science, Office of Biological and Environmental Research (DE-SC0018409), Chemical Sciences, Geoscience and Biosciences Division, Office of Basic Energy Sciences, Office of Science, U.S. Department of Energy (DE-FG02-91ER20021) and MSU AgBioResearch (MICL02598). We thank the Research Technology Support Facility Genomics Core at Michigan State University for the next-generation sequencing.

## Author Contributions

J.K., D.K.K. and F.B. conceived the project, designed experiments and research plan; J.K. and D.K.K performed experiments and data analysis; F.B. supervised the project; J.K and F.B. wrote the manuscript.

---

**Supplementary Figure 1.**
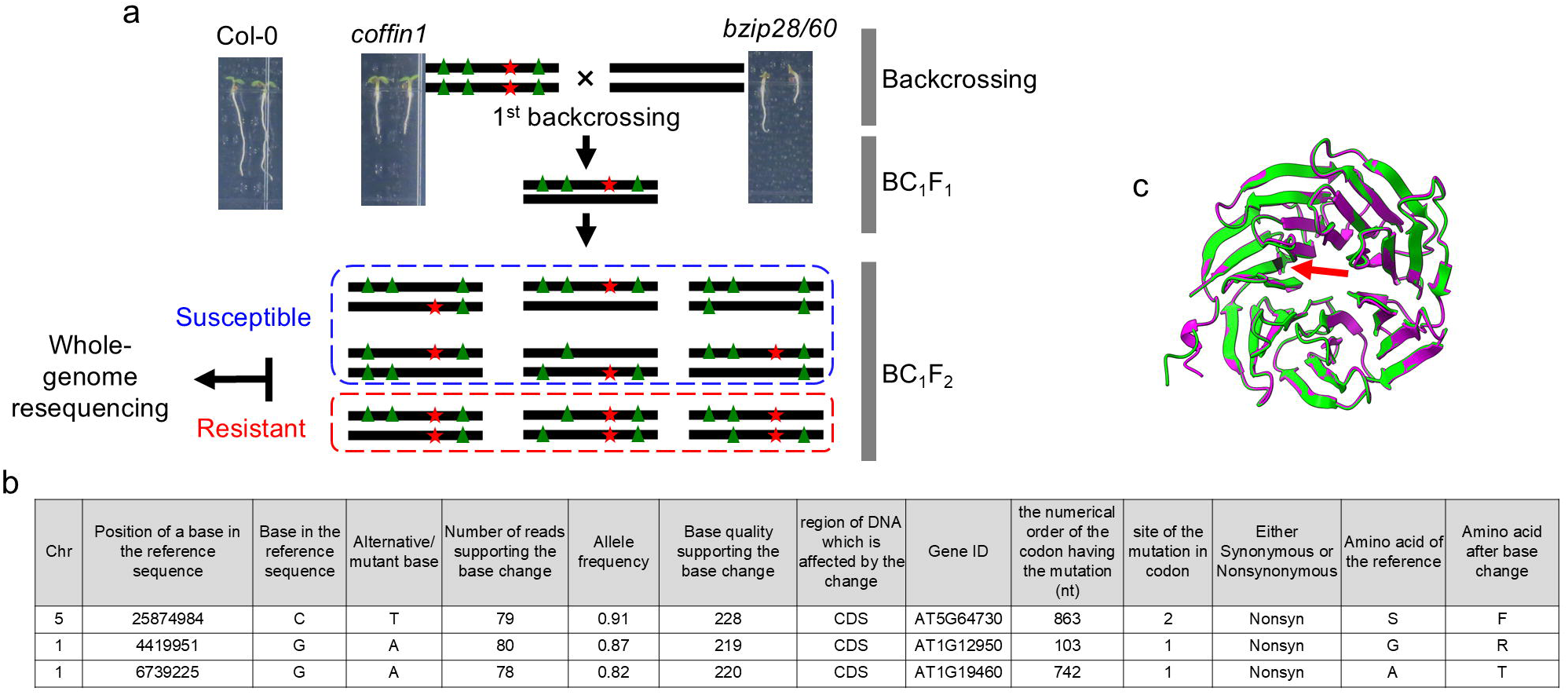
Mapping and identification of the causal mutation in *coffin1*. **a.** Schematic workflow of Bulked Segregant Analysis (BSA) used to identify the coffin1 mutation. The *coffin1* mutant was crossed with *bzip28/60* to generate a BC1F2 mapping population. Genomic DNA was extracted from pools of BC1F2 individuals exhibiting the coffin1 phenotype (TM resistant) and those showing the *bzip28/60* phenotype (TM sensitive) under TM treatment. Whole-genome sequencing was performed on the pooled DNA samples, and single nucleotide polymorphism (SNP) frequencies were analyzed to identify genomic regions linked to the *coffin1* phenotype. **b**. List of the top 3 candidate causal mutations identified by BSA. The *coffin1* mutation, a serine-to-phenylalanine substitution at position 288 (S288F) in the MORG1 gene, is highlighted. **c**. Structural modeling of the MORG1 protein. The wild-type MORG1 protein structure (green ribbon) predicted by AlphaFold3 is superimposed with the predicted structure of the coffin1 mutant protein (S288F, pink ribbon). The red arrow indicates the location of the S288F mutation.

**Supplementary Figure 2.**
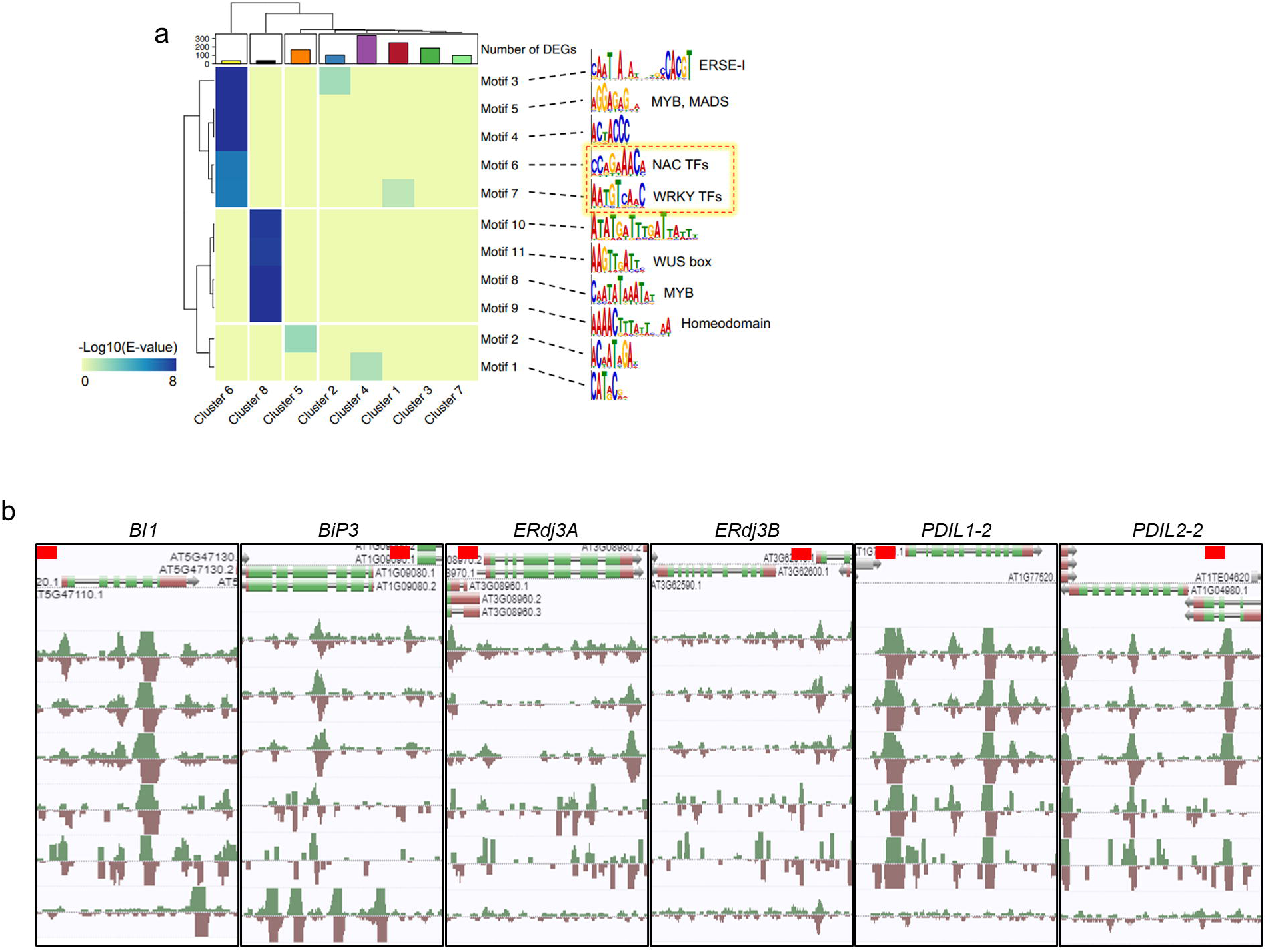
Promoter analysis of DEGs and identification of candidate TFs potentially involved in the partial restoration of UPR gene expression in *coffin1*. **a.** Heatmap showing the enrichment of de novo motifs in the promoters of DEGs. The 1 kb upstream sequences of the transcription start site of each DEG were analyzed using STREME for motif discovery. The heatmap shows the enrichment scores (-log10 p-value) of selected motifs in each of the eight clusters identified by k-means clustering (Figure 3e). Motifs significantly enriched in cluster 6, including NAC and WRKY TF binding sites, are highlighted with box. **b**. Genome browser view of DAP-seq profiles for selected TFs (NAC2, WRKY8, WRKY15, WRKY22, WRKY28, and WRKY45) at the *BI1*, *BiP3*, *ERdj3A*, *ERdj3B*, *PDIL1-2*, and *PDIL2-2* loci. The DAP-seq data were obtained from the Plant Cistrome Database. The y-axis scale (0-15) is consistent across all DAP-seq profiles. The promoter regions of the indicated genes are highlighted with red lines. The enrichment of these TFs in the promoter regions of the indicated genes suggests their potential involvement in regulating UPR gene expression.

**Supplementary Figure 3.**
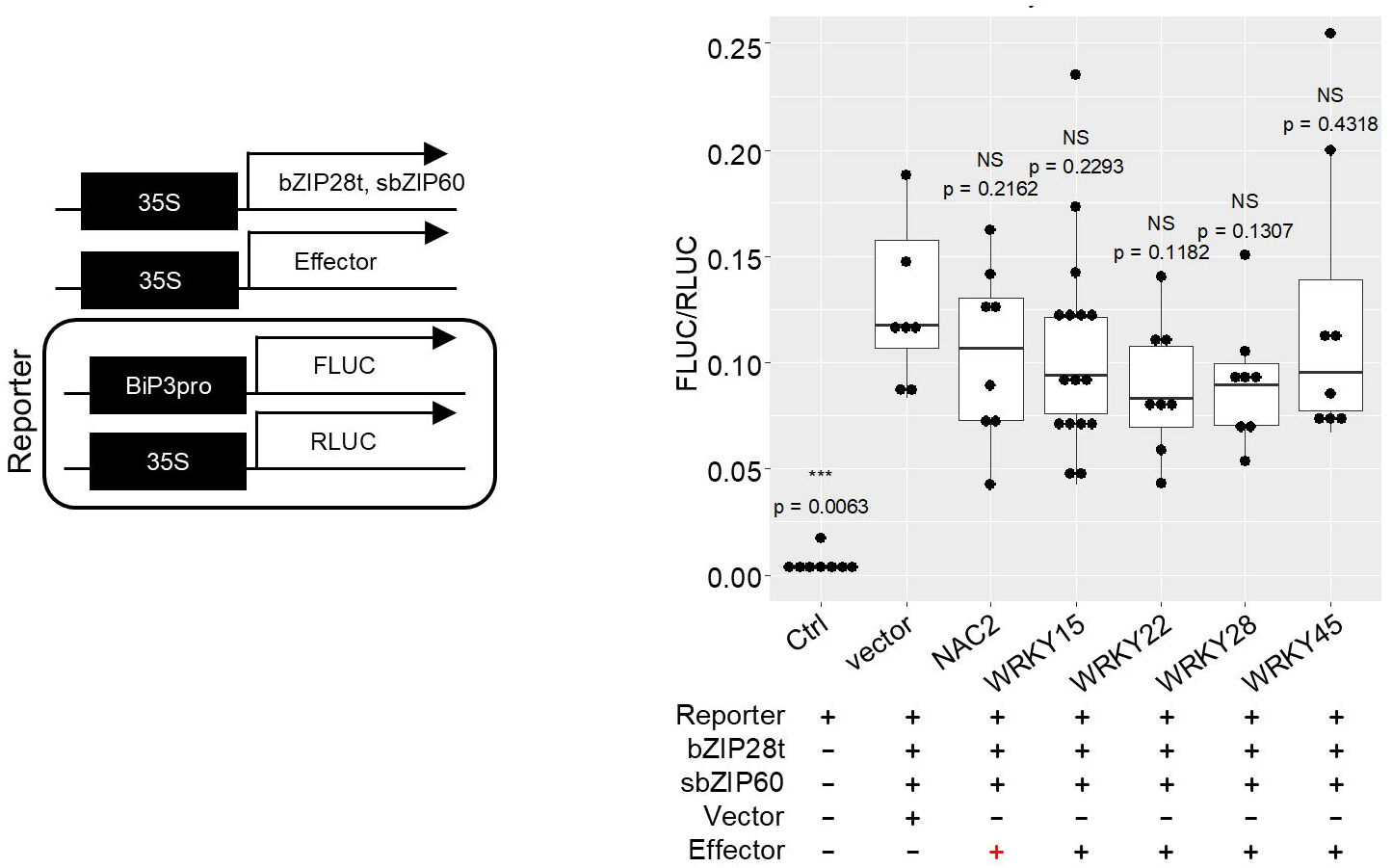
Evaluation of other candidate TFs on BiP3 promoter activity. Dual-luciferase assays were performed in Nicotiana tabacum leaves to assess the effect of the indicated TFs on *BiP3* promoter activity. The reporter construct contains the 1 kb *BiP3* promoter fused to firefly luciferase (FLUC), and the effector constructs express the indicated TFs (NAC2, WRKY15, WRKY22, WRKY28, and WRKY45) under the control of the CaMV 35S promoter. Truncated bZIP28 (bZIP28t) and spliced bZIP60 (sbZIP60) were co-expressed to activate the *BiP3* promoter. Agrobacterium cultures carrying the reporter, effector, and control (empty vector) constructs were infiltrated into N. tabacum leaves as indicated by the plus and minus signs. FLUC activity was normalized to Renilla luciferase (RLUC) activity, which served as an internal control. The normalized FLUC/RLUC values are presented as box plots, with each dot representing an independent biological replicate (n ≥ 7). Statistical significance was determined by Student’s t-test (NS, not significant). None of the tested TFs significantly reduced *BiP3* promoter activity compared to the vector control.

**Supplementary Figure 4.**
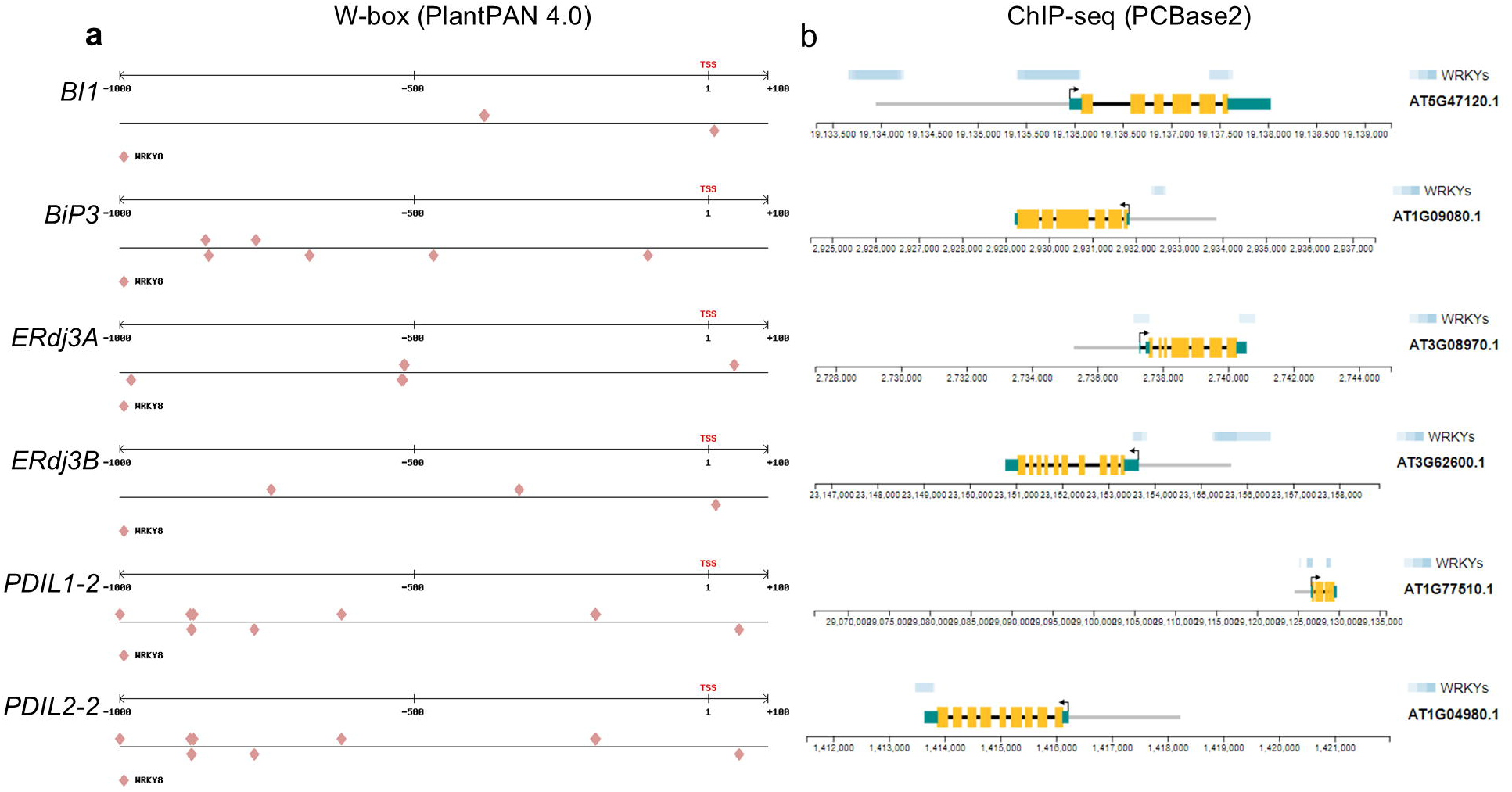
WRKY TFs bind to the promoters of UPR genes. **a**, Schematic representation of the BI1, BiP3, ERdj3A, ERdj3B, PDIL1-2, and PDIL2-2 promoter regions, showing the location of putative W-box elements (TTGACY) within 1 kb upstream to 100 bp downstream of the transcription start site (TSS). The analysis was performed using the PlantPAN 4.0. **b**. ChIP-seq profiles of WRKY TFs at the BI1, BiP3, ERdj3A, ERdj3B, PDIL1-2, and PDIL2-2 loci. The ChIP-seq data, which were generated from a study investigating the role of WRKY TFs in microbe-associated molecular pattern (MAMP)-triggered immunity (MTI) in Arabidopsis (GSE109149), were visualized from the PCBase2.

